# Yap/Taz-Activated Tert-Expressing Acinar Cells Are Required for Pancreatic Regeneration

**DOI:** 10.1101/2021.08.30.458292

**Authors:** Han Na Suh, Moon Jong Kim, Sung Ho Lee, Sohee Jun, Jie Zhang, Randy L. Johnson, Jae-Il Park

**Author notes:** These authors equally contributed to this work (Han Na Suh, Moon Jong Kim).

## Abstract

The expression of TERT (telomerase reverse transcriptase) has been implicated in stem and progenitor cells, which are essential for tissue homeostasis and regeneration. However, the roles of TERT-expressing cells in the pancreas remain elusive. Employing genetically engineered *Tert* knock-in mouse model, herein, we located a rare population of Tert+ acinar cells. While Tert+ cells are quiescent in normal conditions, acinar cell injury leads to mitotic activation of Tert+ cells and subsequent generation of new acinar cells. Moreover, the genetic ablation of Tert+ cells impairs pancreatic regeneration. We further found that Yap/Taz activation is required for the expansion of Tert+ acinar cells. Our results identified Tert+ acinar cells as a distinct subset of acinar cells, which contributes to pancreatic regeneration via Yap/Taz activation.

## INTRODUCTION

Upon injury, the pancreas displays a cellular turnover and proliferation capacity during regeneration [1, 2]. Endocrine pancreas (or also called Langerhans islet) is mainly replenished by self-duplication or dedifferentiation of other cell types in islet [2, 3]. The exocrine pancreas is composed of acinar and ductal epithelial cells and produces the digestive enzymes, including amylases, lipase, and proteinase. Accumulating evidence suggests that a few facultative progenitor cells in the ductal- or acinar compartment play crucial roles in the acinar cell regeneration [4–8]. However, cellular and molecular mechanisms of pancreatic regeneration remain elusive.

TERT, a catalytic subunit of telomerase, is required for telomere maintenance to overcome telomere crisis and cellular senescence [9]. TERT is mainly expressed in self-renewing and regenerating cells, which implies the pivotal roles of TERT-expressing cells in tissue homeostasis and regeneration [10]. A recent study suggested that Tertexpressing (Tert+) cells contribute to liver homeostasis and regeneration [11]. In the intestine, Tert+ cells conditionally divide into epithelial cells and highly proliferative ISCs that are vulnerable to genotoxic stress [12]. Herein, by employing the *Tert* knock-in mouse model [12, 13], we investigated the role of Tert+ cells in pancreatic regeneration.

## RESULTS

### Identification of Tert+ Acinar Cells

To study Tert+ cells in the pancreas, we utilized the *Tert* knock-in mouse model (hereafter referred to as *Tert^TCE/+^*) explicitly expressing tdTomato-CreERT2 (TCE) driven by the endogenous *Tert* promoter [12, 13]. The immunohistochemistry (IHC) analysis located the Tert+ cells in the exocrine pancreas (Figure 1A). Tert+ cells express Amylase and Cpa1, markers for acinar cells (Figures 1B and 1C), but not Ck19, a ductal cell marker (Figure 1E). Fluorescence-activated cell sorting (FACS) analysis showed that in *Tert^TCE/+^* mice, about 0.27± 0.01 % of exocrine pancreatic cells showed the expression of Tert (Figures 1F-1H). Tert+ acinar cells expressed several acinar cell markers, *Amy2a, Cpa1, Ptf1a*, and *Mist*, similar to Tert-acinar cells (Figure 1I). Of note, Tert+ acinar cells also expressed a broad range of other cell type markers, including centroacinar, ductal, and endothelial cells (Figure 1I). However, Tert and Bmi1 doublepositive cells were not located (Figure S1A), implying that Tert+ cells are a somewhat distinct population of Bmi1+ progenitors, previously proposed facultative progenitor cells in the pancreas [6, 8].

**Figure 1.**
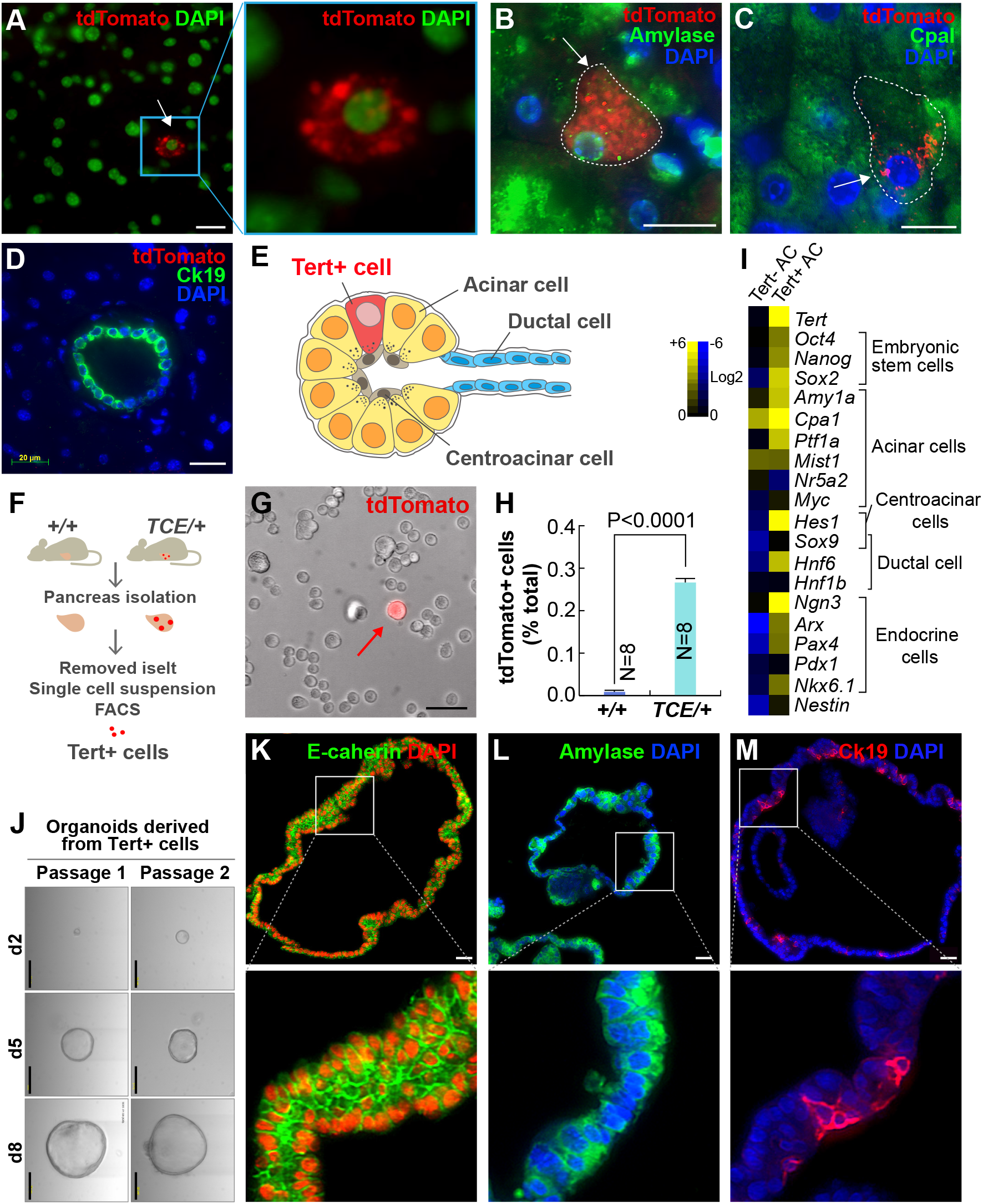
Identification of Tert+ Acinar Cells. (A-E) Identification of Tert+ (Tert-expressing) acinar cells in mouse pancreas. Immunofluorescence (IF) staining of the pancreas of *Tert^TCE/+^* mice for the analyses of tdTomato expression (Tert+ cells) (A). Co-IF staining *Tert^TCE/+^* pancreas for tdTomato (Tert+) and acinar cell markers, Amylase, and CpaI (B and C). Co-IF staining of *Tert^TCE/+^* pancreas for tdTomato (Tert) and Ck19 (ductal cell marker) (D). Scale bars, 20 μm. Illustration of Tert+ cells in mouse pancreas (E). (F-H) Isolation of Tert+ acinar cells. Schematic diagram of Tert+ acinar cell isolation from *Tert^TCE/+^* mice (F). The representative image of isolated Tert+ acinar cell (G); red arrow: Tert+ cell. A single-cell suspension of the exocrine pancreas for FACS and gene expression analysis. Quantification of Tert+ cells isolated from the mouse *Tert^TCE/+^* pancreas by FACS analysis (H); Langerhans islet was removed before single-cell suspension by column. Student’s *t*-test; error bars = SEM; *P < 0.05. (I) Heatmap view of the gene expression pattern of Tert+ and Tert- (Tert not expressing) cells. Tert- and Tert+ cells were analyzed by qRT-PCR (n=3). Approximately five Tert+ and Tert-cells isolated by FACS were subjected to cDNA synthesis and qRT-PCR. Tert-acinar cells prepared from the same *Tert^TCE/+^* mouse pancreatic tissues. (J-M) Tert+ acinar cells form organoids in 3D culture. The representative images of pancreatic organoids derived from Tert+ acinar cells during serial passages (J); d, days. Scale bars, 100 μm. The organoids derived from Tert+ cells display cellular heterogeneity (K-M). IF staining of Tert+ organoids for E-cadherin (epithelial cell marker) (K), Amylase (acinar cell marker) (L), and Ck19 (ductal cell marker) (M); Scale bars, 20 μm. The representative images were displayed.

Next, we tested the proliferation potential of Tert+ acinar cells using in vitro organoid culture system [5, 14]. In three-dimensional (3D) culture, Tert+ acinar cells expanded and developed into organoids until 2 or 3 passages (Figure 1J). However, Tert-acinar cells rarely developed organoids only in the first passage and displayed slow growth, compared to Tert+ cells-derived organoids (Figure S1B). Tert+ cells-derived organoids mainly express the acinar cell marker (Amylase) (Figures 1K-1M).

These results suggest that Tert+ acinar cells belong to a distinct cell population, likely involved in acinar cell replenishment.

### Repopulation of Tert+ Cells by Acinar Cell Injury

Previously, we found that Tert+ intestinal stem cells (ISCs) are quiescent but conditionally activated upon radiation injury [12]. Similarly, Bmi1+ and Doublecortin-like kinase-1 (Dclk1)+ progenitor cells become mitotic after pancreatic injury [5, 6, 8]. To test whether Tert+ acinar cells are engaged in pancreatic regeneration, we administered *Tert^TCE/+^* mice with caerulein (CR). CR is an analog of cholecystokinin, which stimulates excessive digestive enzyme secretion, leading to acute pancreatitis [15]. CR administration induced interstitial edema with intracellular vacuole formation as well as the depletion of acinar cells, followed by the gradual recovery with acinar cell proliferation (Figures S2A-S2C). Interestingly, despite the massive reduction of acinar cells by CR, *Tert* mRNA expression was upregulated in the pancreas of CR-treated mice (Figure S2D). FACS analysis also showed that the number of Tert+ cells was increased (at day 2) and restored to basal level (at day 5) by CR injury (Figures 2A and 2B). While Tert+ cells were non-proliferative in the normal pancreas, Tert+ cells became proliferative upon CR injury, assessed by quantification of Tert+:Ki67+ cells (Figure 2C and 2D). Next, we asked whether mitotically activated Tert+ cells produce acinar cells upon acinar cell injury. To this end, we employed the *Tert^TCE/+^;Rosa26mT/mG* compound strain for the lineage-tracing approach. Tamoxifen (Tam) treatment genetically labels Tert+ cells and their daughter cells with a membranous green fluorescent protein (mGFP) from the membranous Tomato (mTomato) (Figures 2E and 2F). Treatment of *Tert^TCE/+^:Rosa26mT/mG* mice with Tam and subsequent CR treatment showed the mGFP+ acinar cell expansion during pancreatic regeneration (Figures 2G and 2H), indicating that Tert+ cells conditionally repopulate into the newly generated acinar cells upon CR injury. These results suggest that quiescent Tert+ cells conditionally give rise to the acinar cells upon acinar cell injury.

**Figure 2.**
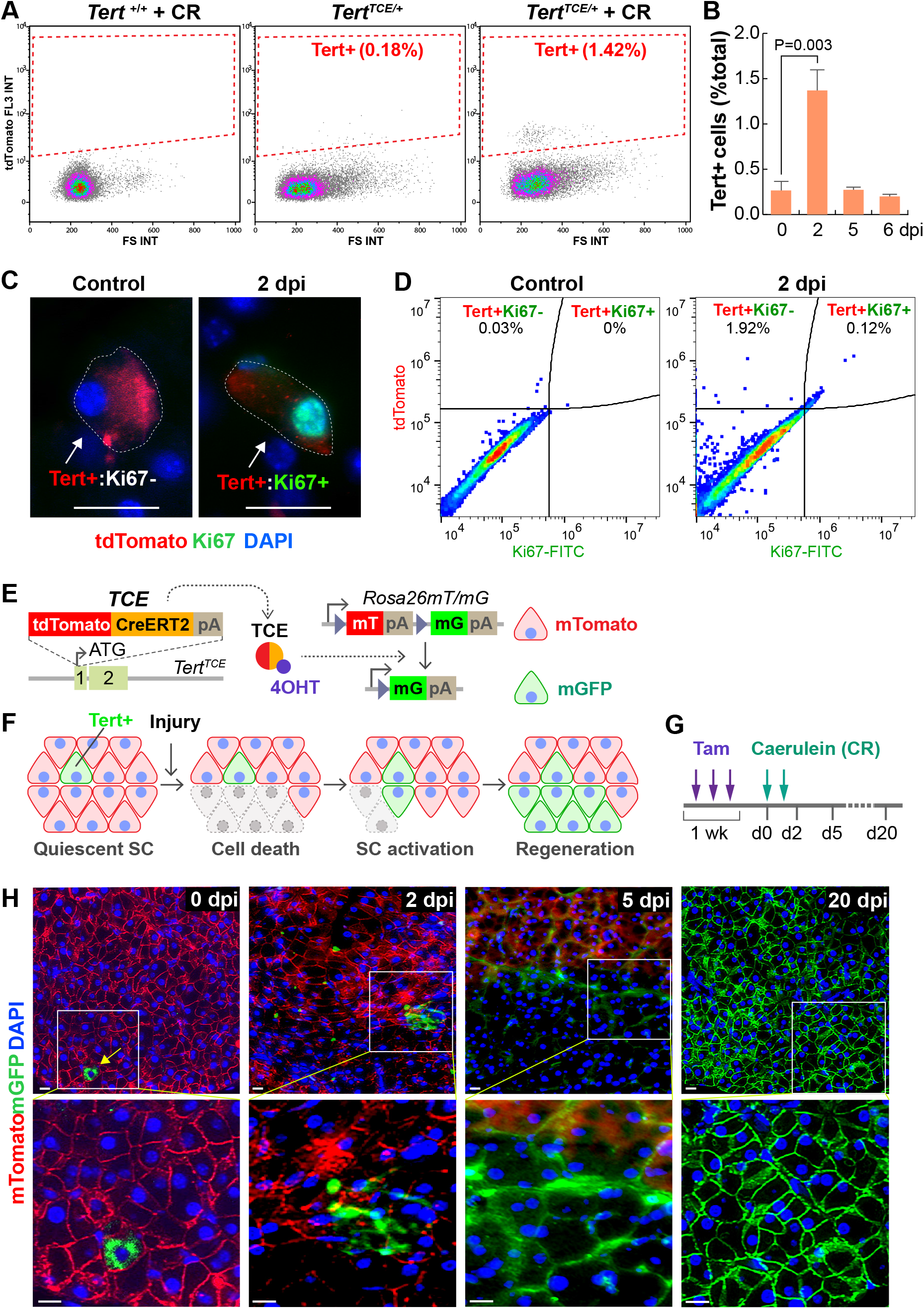
Repopulation of Tert+ Cells by Acinar Cell Injury. (A-B) The increase of the Tert+ cell population after acinar cell injury by CR treatment. FACS analysis of Tert+ cells (tdTomato+) from *Tert^TCE/+^* mice (A). Quantification of Tert+ cells in FACS analysis (B); dpi: days-post injury. (C-D) Mitotic activation of Tert+ Cells after an acinar cell injury. Co-IF staining analysis of mouse *Tert^TCE/+^* pancreas for tdTomato (Tert) and Ki67, a marker for proliferation (C); dpi, days post-injury. Scale bars, 20 μm. FACS analysis of Tert+ and Ki67 doublepositive cells at 2 days after CR treatment (D). (E-H) The lineage-tracing assay of Tert+ cells upon acinar cell injury. Illustration of lineage-tracing of Tert+ cells (*Tert^TCE/+^; Rosa26mT/mG*). Tam administration activates tdTomato-CreERT2 (TCE), which leads to the expression of membranous GFP (mGFP) in the progenies of Tert+ (E and F). Scheme of the lineage tracing assay (G); *Tert^TCE/+^;Rosa26mT/mG* mice were administrated with Tam (1 week) and CR. Microscopic analysis of Tert+ cell lineage. At 0, 2, 5, 20 dpi, cryo-sectioned pancreatic samples were analyzed for mGFP and mTomato expression (H). Scale bars, 20 μm. The representative images were displayed.

### Impaired Pancreatic Regeneration by Conditional Ablation of Tert+ Cells

Having determined the quiescence exit and repopulation of Tert+ cells upon acinar cell injury, we next asked whether Tert+ cells are required for pancreatic regeneration using cell ablation assays. Upon Tam administration, *Tert^TCE/+^;Rosa26DTA* compound strain expresses diphtheria toxin A (DTA), specifically in Tert+ cells, which results in selective depletion/death of Tert+ cells (Figures 3A and 3B) [12]. Tert+ cell-depleted mice (Tam-treated) were then under CR injury for evaluation of pancreatic regeneration. While vehicle and CR-treated control mice showed the completion of pancreatic regeneration at 8 dpi, Tert+ cell-depleted mice (Tam and CR-treated) displayed the impaired pancreatic regeneration, represented by the failure in the acinar cell (Amylase+) regeneration (Figures 3C and 3D). Of note, upon acinar cell injury, more Ck19+ cells were observed in the Tert+ cell ablated mice than control mice (Figure 3E). These results suggest that Tert+ acinar cells are required for pancreatic regeneration.

**Figure 3.**
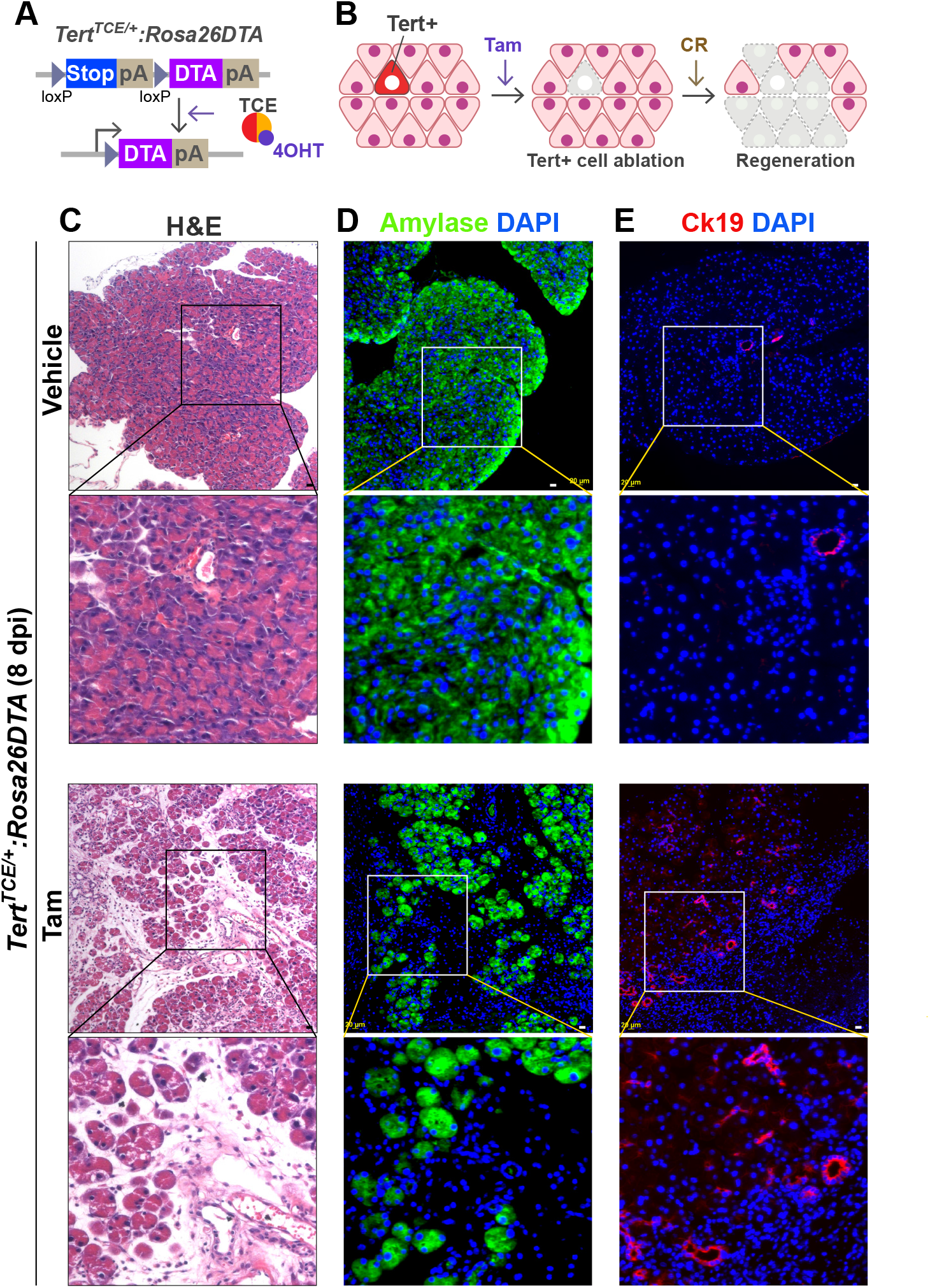
Impaired Pancreatic Regeneration by Conditional Ablation of Tert+ Cells. (A) Illustration of conditional ablation of Tert+ cells (*Tert^TCE/+^;Rosa26DTA*). Tam administration activates TCE and subsequently induces the expression of DTA specifically in Tert+ cells, resulting in the ablation of Tert+ cells. (C-E) Impairment of pancreatic regeneration by Tert+ cell ablation. H&E staining analysis of mouse pancreas (C). IF staining for Amylase (D) and Ck19 (E). Scale bars, 20 μm. The representative images were displayed.

### Hippo Signaling is Required for Tert+ cell Expansion during Pancreatic Regeneration

To gain mechanistic insight into how Tert+ cells contribute to pancreatic regeneration, we performed gene expression analysis of CR-treated pancreas. Among several developmental pathways, Hippo downstream target genes (*Ctgf, Ankrd1*, and *Cyr61*) were significantly upregulated upon CR (1 dpi) (Figure S3A). Hippo signaling regulates the self-renewal and expansion of stem and progenitor cells [16–18] and plays crucial roles in tissue regeneration of intestine [19], heart [20], and liver [21]. Moreover, the Hippo signaling contributes to pancreatic development [22, 23], which led us to test whether Hippo signaling is required for the expansion of Tert+ cells using *Yap* and *Taz* conditional knock-out (CKO) mouse models. We genetically ablated *Yap/Taz* alleles in Tert+ cells by Tam administration of *Tert^TCE/+^;Yap^fl/fl^:Taz^fl/fl^* mice (Figure 4A). Then, we determined the impacts of *Yap/Taz* CKO in Tert+ cells on CR-induced pancreatic regeneration. Indeed, *Tert^TCE/+^;Yap^Δ/Δ^;Taz^Δ/Δ^* mice showed the impairment of pancreatic regeneration along with reduced acinar cell generation and relatively increased ductal cell marker, similar to Tert+ cell ablation phenotype (Figures 4B-4D). In vitro culture of Tert+ cells-derived organoids, also showed the growth retardation by *Yap/Taz* CKO compared to controls (Figures 4E and 4F). These results suggest that Yap/Taz activation is cell-autonomously required for the Tert+ cells-mediated pancreatic regeneration (Figure 4G).

**Figure 4.**
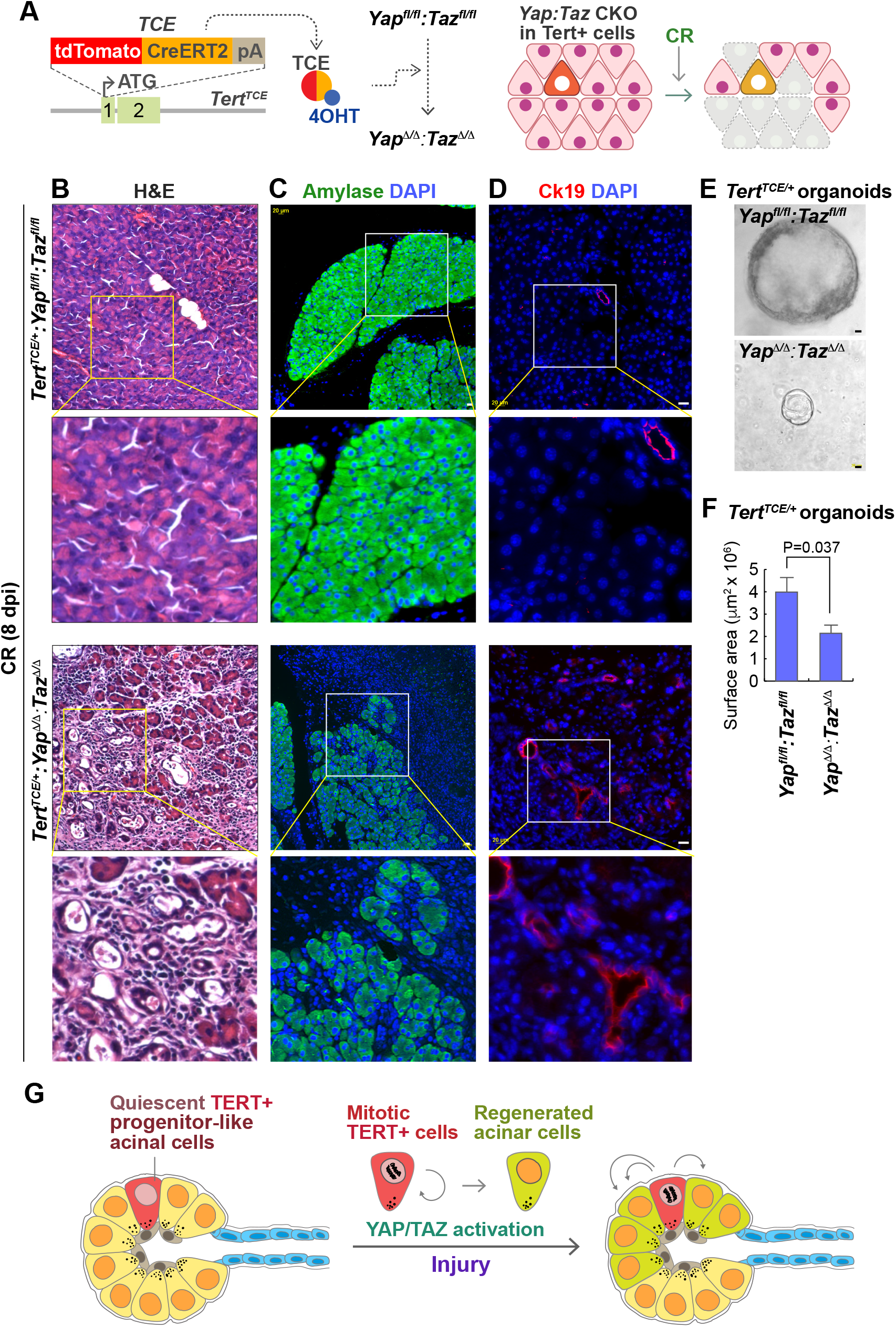
Yap/Taz Activation in Tert+ cells Is Required for Pancreatic Regeneration. (A) Illustration of *Yap/Taz* CKO in Tert+ cells (*Tert^TCE/+^;Yap^fl/fl^;Taz^fl/fl^*) (B-D) Impairment of pancreatic regeneration by *Yap/Taz* CKO in Tert+ cells. H&E staining analysis of mouse pancreatic tissue samples (B). IF staining of mouse pancreas for Amylase (C) and Ck19 (D). Scale bars, 20 μm. (E-F) Reduced organoid growth by *Yap/Taz* CKO in Tert+ cells. Representative organoids images (E). Quantification of organoid surface area (F). The area was analyzed by AxioVision. Student’s *t*-test; error bars = SEM. (G) Model of Tert+ acinar cell expansion during exocrine pancreatic regeneration. Upon acinar cell injury, non-proliferative Tert+ acinar cells become mitotic and generate the daughter cells. Intrinsic YAP/TAZ activation in Tert+ acinar cells is required for pancreatic regeneration. The representative images were displayed.

## DISCUSSION

Despite the roles of facultative pancreatic progenitor cells in pancreatic regeneration [2, 5, 6, 8, 14], the mechanisms of pancreatic regeneration remain ambiguous. Tert+ cells were recently proposed as the self-renewing tissue stem cells contributing to intestinal and liver regeneration [11, 12]. Here, employing the *Tert* knock-in mouse model, we located Tert+ acinar cells, which contribute to pancreatic regeneration.

The current concept in mouse exocrine regeneration is that a rare population of progenitor cells drives the acinar ductal regeneration. For instance, the progenitor cell-like acinar cells expressing Bmi1 or Dclk1 are engaged in pancreatic regeneration [5, 8]. The role of Tert+ cells in pancreatic regeneration seems similar to that of Bmi1+ or Dclk1+ progenitor cells. Because upon acinar cell injury, Tert+ cells acutely repopulate into acinar cells (see Figure 2D). Nonetheless, Tert+ cells do not express Bmi1 (see Figure S1A). Also, Tert+ cells are relatively rare than Bmi1+ cells (Tert+ cells ≅ 0.49%; Bmi1+ cells ≅ 2.5%) [6], suggesting that Tert+ cells are somewhat distinct from Bmi1 + cells. In line with this, despite similar roles of both Tert+ and Bmi1 ISCs during regeneration, Tert+ and Bmi1 ISCs co-exist as distinct cell populations [12, 24]. It should also be noted that, though Tert+ cell ablation impairs the pancreatic regeneration, our results cannot completely rule out the compensation of Tert+ cell loss by other progenitor cells in pancreatic reconstitution and cell differentiation. Moreover, potential cell plasticity conditionally generating Tert+ cells should be considered [25, 26]. These questions need to be addressed by comparative transcriptional profiling of Tert+, Bmi1+, and Dclk1+ cells as well as single-cell RNA-seq.

Cell-lineage tracing has identified various progenitor cells involved in tissue homeostasis and regeneration. However, it is still unclear how the progenitor cells are activated and expanded. The Hippo pathway is evolutionarily conserved and controls organ size by phosphorylating-mediated inhibition of YAP and TAZ [27, 28]. By genetic targeting of *Yap* and *Taz* alleles in Tert+ cells, we found that Yap/Taz activation is indispensable for Tert+ cell expansion for pancreatic regeneration (see Figure 4). The loss of cell-cell contact, cell adhesion, or cytoskeletal integrity leads to transient Hippo/MST inhibition and subsequent YAP/TAZ activation [29]. CR-induced acute pancreatitis is accompanied by the loss of cell-cell contact and disruption of cell adhesion. Thus, it is plausible that the relief of mechanical stress by acinar cell integrity loss might trigger the activation of endogenous Yap/Taz, which then activates Tert+ cells for the initiation of pancreatic regeneration.

Together, our study identified of Tert+ acinar cells as a distinct subset of acinar cells, which are required for pancreatic regeneration via Yap/Taz-mediated Tert+ cell activation.

## SUPPLEMENTAL INFORMATION

Supplemental Information includes three figures and two tables.

## AUTHOR CONTRIBUTIONS

H.N.S., S.H.L., and J.-I.P. conceived the experiments. R.L.J provided *Yap^fl/fl^:Taz^fl/fl^* mice. H.N.S., S.H.L., M.J.K., S.J., J.Z., and J.-I.P. performed the experiments. H.N.S., M.J.K., S.H.L., and J.-I.P. analyzed the data. M.J.K., H.N.S., and J.-I.P. wrote the manuscript.

## ACKNOWLEDGMENTS

We thank Christopher Cervantes, Esther Lien, and Young Sun Oh for technical assistance. This work was supported by the Duncan Family Institute Research Program, the University Cancer Foundation (IRG-08-061-01), the Center for Stem Cell and Developmental Biology (MD Anderson Cancer Center), an Institutional Research Grant (MD Anderson Cancer Center), a New Faculty Award (CA016672), a Metastasis Research Center Grant (MD Anderson Cancer Center), and the Uterine SPORE Career Enhancement Program (MD Anderson Cancer Center).

## METHODS

### Mouse experiments

*Tert^TCE/+^* mice [13] and *Yap^fl/fl^:Taz^fl/fl^* mice [20, 30] were utilized as previously described. *Gt(ROSA)26Sor^tm4(ACTB-tdTomato,-EGFP)Luo^/J* (Jax strain 007576) and *Gt(ROSA)26Sor^tm1(DTA)Lky^/J* (Jax strain 009669) were purchased from Jackson Laboratory. Mice (older than six weeks) were injected with Tamoxifen (Tam; Sigma) for cell lineage-tracing, cell ablation, or CKO of *Yap/Taz* in Tert+ cells. Tamoxifen was dissolved in corn oil (Fisher) at a final concentration of 10 mg/ml for intraperitoneal administration (50 mg/kg), as previously performed [12]. All mice were maintained at the MD Anderson animal facility with a 12 h light/dark schedule and food and water supply and treated in compliance with the Institutional Animal Care and Use Committee guideline of MD Anderson Cancer Center.

### Tert+ acinar cell isolation and FACS analysis

For Tert+ acinar cell isolation experiment, the pancreas was digested with 2.5 mg/ml Collagenase D (Roche 1108866001), supplemented with 0.1 mg/ml DNAse I (Sigma DN25) for 40 min and collected through 70 μm (BD 087712) cell strainers for removing islets of Langerhans. Purified cells were suspended into the single cells by TrypLE (Life Technologies 12605). Cells were then resuspended in PBS with 10% fetal bovine serum (FBS) and stained with SYTOX^®^ Blue (Life Technologies S34857) to exclude dead cells. Based on tdTomato fluorescence, Tert+ acinar cells were sorted into Tert+ and Tertcells among live cells based on the gate and collected by FACS (MoFlo^®^Astrios™, Beckman Coulter). C57BL/6J mice were used as negative controls for gating. Sorted Tert+ and Tert-cells were subsequently analyzed for the gene expression profiling or organoid culture. T quantify the proliferative potential of Tert+ cells, sorted Tert+ cells were fixed with 4% paraformaldehyde, blocked, incubated with FITC-conjugated Ki67 antibody and stained with DAPI. The population of Tert+:Ki67+ was assessed by FACS (Gallios™ 561, Beckman Coulter).

### Gene expression analysis

Isolated Tert+ and Tert-cells (≤ 5 cells) were subjected to synthesize the complementary DNA (cDNA) using REPLI-g WTA Single Cell Kit (QIAGEN) and analyzed for gene expression by qRT-PCR. 18S ribosomal RNA (*18S rRNA*) was used as an endogenous control for normalization. All qRT-PCR experiments were performed using intron-spanning primers. Fold induction was quantified using a 2^-ΔΔCT^ method. For Figure 1I, ΔΔCT values were displayed using heatmap software (http://bar.utoronto.ca/). Primer sequences are listed in Table S1.

### Tert+ organoid culture

FACS sorted Tert- or Tert+ acinar cells were resuspended in Matrigel (BD 356231) for seeding. After incubation for 5 min at 37°C to solidify the Matrigel, cells were grown in the modified pancreatic organoid culture medium [5, 14]; supplemented with Advanced DMEM/F-12 (Life Technologies 12634-010) supplemented with 1% penicillin/streptomycin, 1% GlutaMAX (Life Technologies 35050061), 10 mM HEPES (Life Technologies, 2% (v/v) B27 supplement (Life Technologies 12587-010), 1 mM N-Acetylcysteine (MP Biomedicals ICN19460305), 0.5 μg/ml R-spondin (R&D 3474-RS), 10 mM nicotinamide (Sigma N0636), 10 nM recombinant human [Leu15]-gastrin I (Sigma G9145), 50 ng/ml recombinant mouse EGF (Peprotech, 315-09), 100 ng/ml recombinant human FGF10 (Peprotech 100-26), 25 ng/ml recombinant human Noggin (Peprotech 250-38), 10 μM Y-27632 (Cayman Chemical 10005583), and 1mM Jagged-1 (AnaSpec AS-61298). Organoids were maintained at 37°C, and 5% carbon dioxide atmosphere and the culture medium was changed every other day. Organoid imaging and measurement were performed using a Zeiss Z1 microscope with AxioVision software (Zeiss).

### Acute pancreatitis mouse model

Mice weighing 20 to 25 g were used. Before the experiments, mice were fasted for 12 h and allowed water ad libitum. In the caerulein (CR; Sigma) treated group, mice were injected intraperitoneally with CR (400 μg/kg) dissolved in phosphate-buffered saline (PBS; BD) at a 1 h interval six times for one day.

### Histology and immunohistochemistry (IHC)

The pancreatic samples were fixed in 10% neutral buffered formalin overnight and embedded in paraffin. Tissue samples were then sectioned (5 μm), and H&E staining was performed following standard procedure. For IHC, slides were deparaffinized, rehydrated, processed for antigen retrieval, blocked, incubated with the primary antibody, and fluorescence or peroxidase-conjugated secondary antibody. Next, slides were mounted with DAPI (Invitrogen P36935), sealed, and photographed using a microscope. For comparison among the experiment groups, images were captured with the same exposure time. All antibody information is listed in Table S2.

### Statistical analyses

The Student’s *t*-test was used for comparisons of two samples. P values < 0.05 were considered significant. Error bars indicate standard error (SEM). The number of biological and experimental replicates is ≥ 3, otherwise mentioned in Figure Legends.

## Supplementary Information

### Supplementary Figure Legends

**Figure S1.**
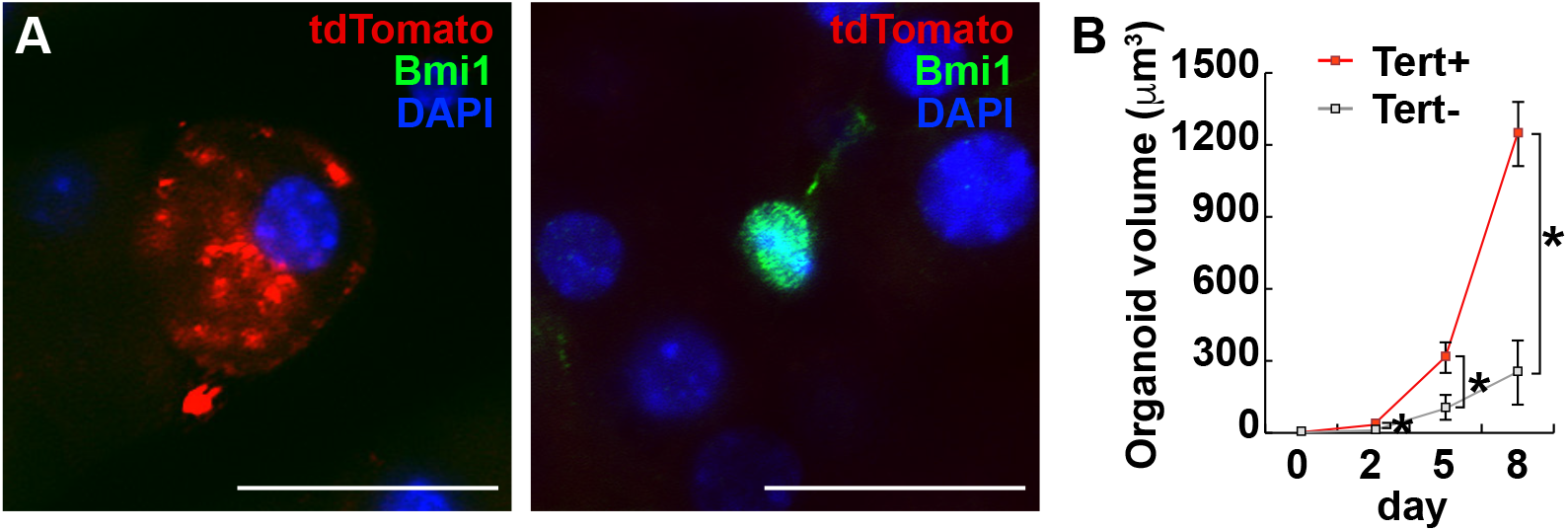
Characterization of Tert+ Cells in Pancreas (related to Figure 1) (A) Co-IF staining of mouse *Tert^TCE/+^* pancreas for tdTomato (Tert) and Bmi1 (A). At least five 200 × magnified fields from at least 3 mice were analyzed. Scale bars, 20 μm. (B) Growth of Tert+ cells-derived organoids in 1^st^ passages. Of note, Tert-cells rarely formed organoids and could not be maintained next passage. Only growing Tertorganoids were analyzed and displayed.

**Figure S2.**
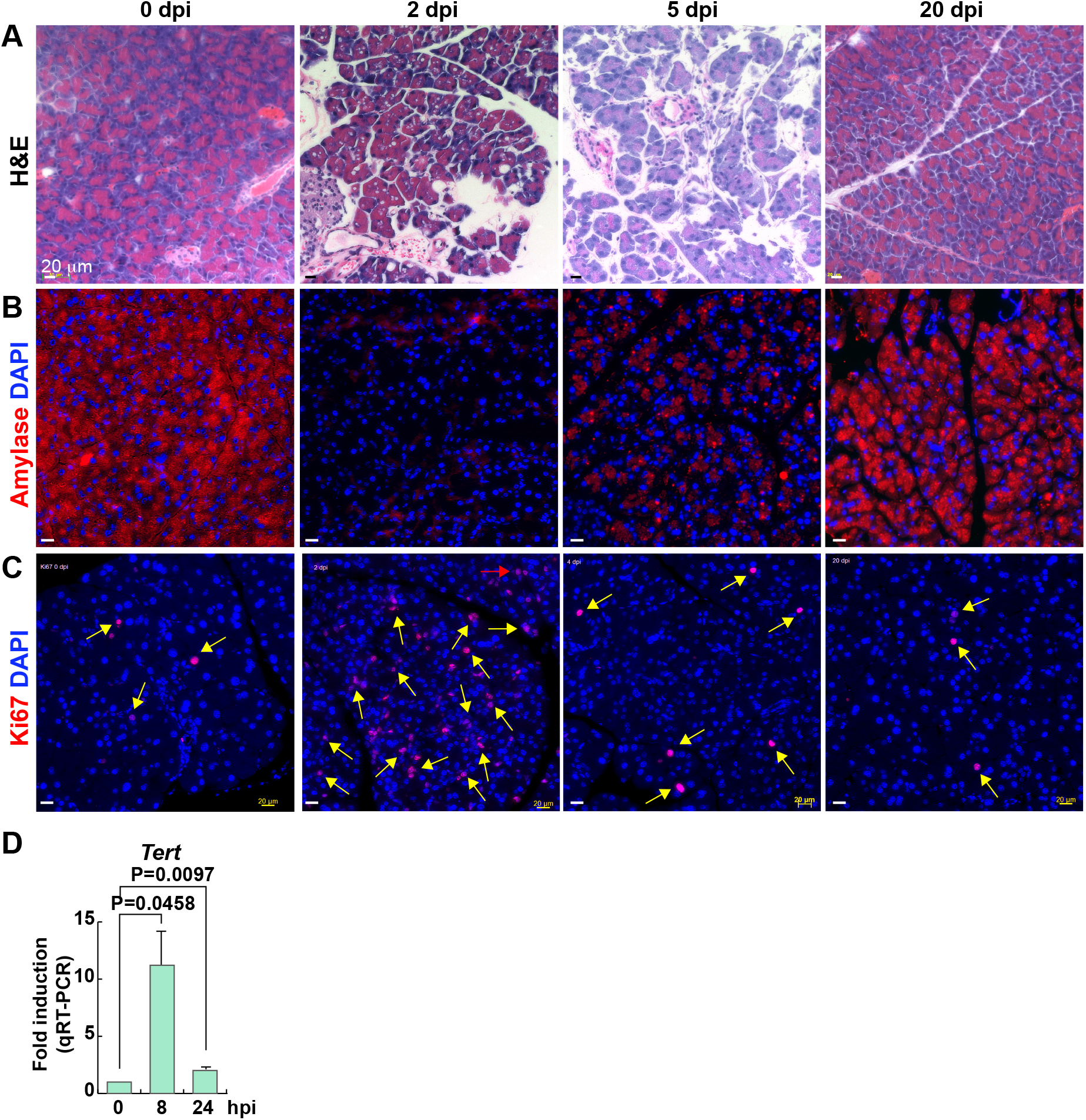
Pancreatic Regeneration after Acute Pancreatitis (related to Figure 2) (A-C) Pancreatic regeneration after CR treatment. dpi, days post-injury. H&E staining (A). IF staining of mouse pancreas for Amylase (B) and Ki67 (C). Scale bars, 20 μm. (D) Upregulation of *Tert* mRNA in the pancreas after CR treatment. hpi, hours postinjury.

**Figure S3.**
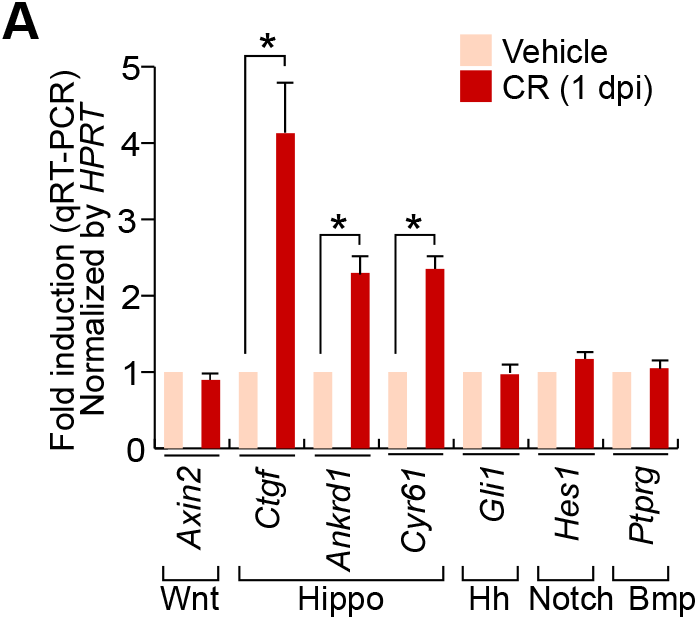
Activation of Hippo Downstream Target Genes after CR Treatment (related to Figure 4) (A) Gene expression analysis of the regenerating pancreas; CR (1 dpi). After CR treatment, the mouse whole pancreas samples were analyzed by qRT-PCR. dpi, days post-injury. Student’s *t*-test; error bars = SEM; *P < 0.05.

### Supplementary Tables

**Table S1.**
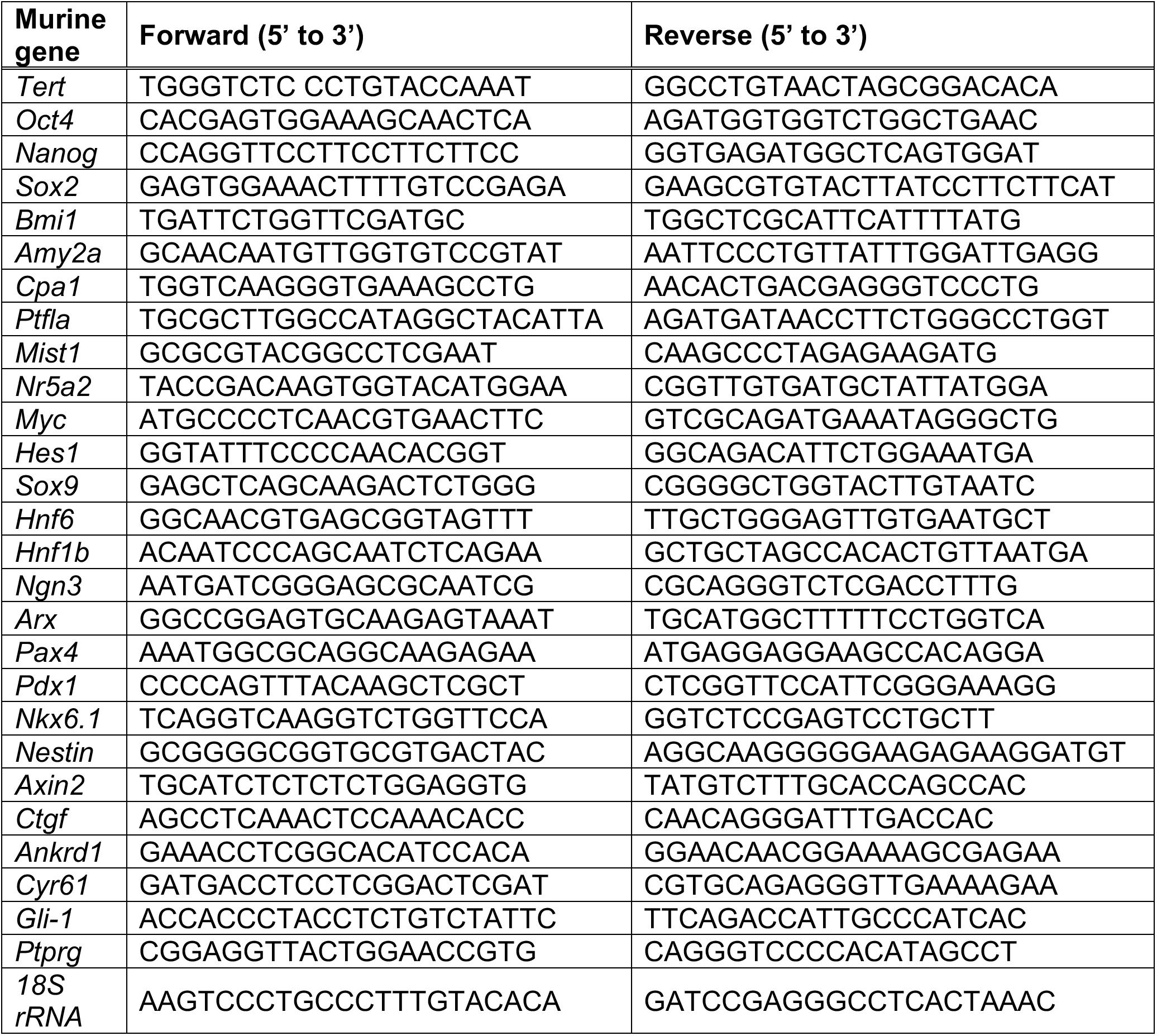
Primer Sequence Information (related to Figures 1 and S3) List of all primer sequences used for qRT-PCR

**Table S2.** Antibody Information for IHC or IF (related to Figures 1-4, S1, and S2)

| **Protein** | **Company (cat #)** | **Dilution** |
| --- | --- | --- |
| Amylase | Abcam (ab21156) | 1/200 |
| Cpa1 | R&D (AF2765) | 1/100 |
| Ck19 | Abcam (ab133496) | 1/200 |
| Bmi1 | Abcam (ab14389) | 1/100 |
| RFP | Invitrogen (MA5-15257) | 1/100 |
| Ki67 | Abcam (ab16667) | 1/100 |
| E-cadherin | BD (610182) | 1/200 |
| Yap | Abcam (ab52771) | 1/100 |
| Ctgf | Abcam (ab6992) | 1/100 |
| Ki67-FITC | eBioscience (11-5698-82) | 1/50 |

## REFERENCES

1. Murtaugh LC, Keefe MD. Regeneration and repair of the exocrine pancreas. Annu Rev Physiol. 2015;77:229–49. Epub 2014/11/12. doi: 10.1146/annurev-physiol-021014-071727. PubMed PMID: 25386992; PubMed Central PMCID: PMCPMC4324082.

2. Zhou Q, Melton DA. Pancreas regeneration. Nature. 2018;557(7705):351–8. Epub 2018/05/18. doi: 10.1038/s41586-018-0088-0. PubMed PMID: 29769672; PubMed Central PMCID: PMCPMC6168194.

3. Xu X, D’Hoker J, Stange G, Bonne S, De Leu N, Xiao X, et al. Beta cells can be generated from endogenous progenitors in injured adult mouse pancreas. Cell. 2008;132(2):197–207. Epub 2008/02/05. doi: 10.1016/j.cell.2007.12.015. PubMed PMID: 18243096.

4. Strobel O, Dor Y, Alsina J, Stirman A, Lauwers G, Trainor A, et al. In vivo lineage tracing defines the role of acinar-to-ductal transdifferentiation in inflammatory ductal metaplasia. Gastroenterology. 2007;133(6):1999–2009. Epub 2007/12/07. doi: 10.1053/j.gastro.2007.09.009. PubMed PMID: 18054571; PubMed Central PMCID: PMCPMC2254582.

5. Westphalen CB, Takemoto Y, Tanaka T, Macchini M, Jiang Z, Renz BW, et al. Dclk1 Defines Quiescent Pancreatic Progenitors that Promote Injury-Induced Regeneration and Tumorigenesis. Cell Stem Cell. 2016;18(4):441–55. Epub 2016/04/09. doi: 10.1016/j.stem.2016.03.016. PubMed PMID: 27058937; PubMed Central PMCID: PMCPMC4826481.

6. Sangiorgi E, Capecchi MR. Bmi1 lineage tracing identifies a self-renewing pancreatic acinar cell subpopulation capable of maintaining pancreatic organ homeostasis. Proc Natl Acad Sci U S A. 2009;106(17):7101–6. Epub 2009/04/18. doi: 10.1073/pnas.0902508106. PubMed PMID: 19372370; PubMed Central PMCID: PMCPMC2678421.

7. Desai BM, Oliver-Krasinski J, De Leon DD, Farzad C, Hong N, Leach SD, et al. Preexisting pancreatic acinar cells contribute to acinar cell, but not islet beta cell, regeneration. J Clin Invest. 2007;117(4):971–7. Epub 2007/04/04. doi: 10.1172/JCI29988. PubMed PMID: 17404620; PubMed Central PMCID: PMCPMC1838936.

8. Fukuda A, Morris JPt, Hebrok M. Bmi1 is required for regeneration of the exocrine pancreas in mice. Gastroenterology. 2012;143(3):821–31 e2. Epub 2012/05/23. doi: 10.1053/j.gastro.2012.05.009. PubMed PMID: 22609312; PubMed Central PMCID: PMCPMC3485080.

9. Greider CW, Blackburn EH. Identification of a specific telomere terminal transferase activity in Tetrahymena extracts. Cell. 1985;43(2 Pt 1):405–13. Epub 1985/12/01. doi: 10.1016/0092-8674(85)90170-9. PubMed PMID: 3907856.

10. Gunes C, Rudolph KL. The role of telomeres in stem cells and cancer. Cell. 2013;152(3):390–3. Epub 2013/02/05. doi: 10.1016/j.cell.2013.01.010. PubMed PMID: 23374336.

11. Lin S, Nascimento EM, Gajera CR, Chen L, Neuhofer P, Garbuzov A, et al. Distributed hepatocytes expressing telomerase repopulate the liver in homeostasis and injury. Nature. 2018;556(7700):244–8. Epub 2018/04/06. doi: 10.1038/s41586-018-0004-7. PubMed PMID: 29618815; PubMed Central PMCID: PMCPMC5895494.

12. Suh HN, Kim MJ, Jung YS, Lien EM, Jun S, Park JI. Quiescence Exit of Tert(+) Stem Cells by Wnt/beta-Catenin Is Indispensable for Intestinal Regeneration. Cell Rep. 2017;21(9):2571–84. Epub 2017/12/01. doi: 10.1016/j.celrep.2017.10.118. PubMed PMID: 29186692; PubMed Central PMCID: PMCPMC5726811.

13. Jun S, Jung YS, Suh HN, Wang W, Kim MJ, Oh YS, et al. LIG4 mediates Wnt signalling-induced radioresistance. Nat Commun. 2016;7:10994. Epub 2016/03/25. doi: 10.1038/ncomms10994. PubMed PMID: 27009971; PubMed Central PMCID: PMCPMC4820809.

14. Wollny D, Zhao S, Everlien I, Lun X, Brunken J, Brune D, et al. Single-Cell Analysis Uncovers Clonal Acinar Cell Heterogeneity in the Adult Pancreas. Dev Cell. 2016;39(3):289–301. Epub 2016/12/08. doi: 10.1016/j.devcel.2016.10.002. PubMed PMID: 27923766.

15. Elsasser HP, Adler G, Kern HF. Time course and cellular source of pancreatic regeneration following acute pancreatitis in the rat. Pancreas. 1986;1(5):421–9. Epub 1986/01/01. PubMed PMID: 2436216.

16. Lian I, Kim J, Okazawa H, Zhao J, Zhao B, Yu J, et al. The role of YAP transcription coactivator in regulating stem cell self-renewal and differentiation. Genes Dev. 2010;24(11):1106–18. Epub 2010/06/03. doi: 10.1101/gad.1903310. PubMed PMID: 20516196; PubMed Central PMCID: PMCPMC2878649.

17. Camargo FD, Gokhale S, Johnnidis JB, Fu D, Bell GW, Jaenisch R, et al. YAP1 increases organ size and expands undifferentiated progenitor cells. Curr Biol. 2007;17(23):2054–60. Epub 2007/11/06. doi: 10.1016/j.cub.2007.10.039. PubMed PMID: 17980593.

18. Cao X, Pfaff SL, Gage FH. YAP regulates neural progenitor cell number via the TEA domain transcription factor. Genes Dev. 2008;22(23):3320–34. Epub 2008/11/19. doi: 10.1101/gad.1726608. PubMed PMID: 19015275; PubMed Central PMCID: PMCPMC2600760.

19. Cai J, Zhang N, Zheng Y, de Wilde RF, Maitra A, Pan D. The Hippo signaling pathway restricts the oncogenic potential of an intestinal regeneration program. Genes Dev. 2010;24(21):2383–8. Epub 2010/11/03. doi: 10.1101/gad.1978810. PubMed PMID: 21041407; PubMed Central PMCID: PMCPMC2964748.

20. Xin M, Kim Y, Sutherland LB, Murakami M, Qi X, McAnally J, et al. Hippo pathway effector Yap promotes cardiac regeneration. Proc Natl Acad Sci U S A. 2013;110(34):13839–44. Epub 2013/08/07. doi: 10.1073/pnas.1313192110. PubMed PMID: 23918388; PubMed Central PMCID: PMCPMC3752208.

21. Loforese G, Malinka T, Keogh A, Baier F, Simillion C, Montani M, et al. Impaired liver regeneration in aged mice can be rescued by silencing Hippo core kinases MST1 and MST2. EMBO Mol Med. 2017;9(1):46–60. Epub 2016/12/13. doi: 10.15252/emmm.201506089. PubMed PMID: 27940445; PubMed Central PMCID: PMCPMC5210079.

22. Ardestani A, Maedler K. The Hippo Signaling Pathway in Pancreatic beta-Cells: Functions and Regulations. Endocr Rev. 2018;39(1):21–35. Epub 2017/10/21. doi: 10.1210/er.2017-00167. PubMed PMID: 29053790.

23. Gao T, Zhou D, Yang C, Singh T, Penzo-Mendez A, Maddipati R, et al. Hippo signaling regulates differentiation and maintenance in the exocrine pancreas. Gastroenterology. 2013;144(7):1543–53, 53 e1. Epub 2013/03/05. doi: 10.1053/j.gastro.2013.02.037. PubMed PMID: 23454691; PubMed Central PMCID: PMCPMC3665616.

24. Sangiorgi E, Capecchi MR. Bmi1 is expressed in vivo in intestinal stem cells. Nat Genet. 2008;40(7):915–20. Epub 2008/06/10. doi: 10.1038/ng.165. PubMed PMID: 18536716; PubMed Central PMCID: PMCPMC2906135.

25. Puri S, Folias AE, Hebrok M. Plasticity and dedifferentiation within the pancreas: development, homeostasis, and disease. Cell Stem Cell. 2015;16(1):18–31. Epub 2014/12/04. doi: 10.1016/j.stem.2014.11.001. PubMed PMID: 25465113; PubMed Central PMCID: PMCPMC4289422.

26. Kopp JL, Grompe M, Sander M. Stem cells versus plasticity in liver and pancreas regeneration. Nat Cell Biol. 2016;18(3):238–45. Epub 2016/02/26. doi: 10.1038/ncb3309. PubMed PMID: 26911907.

27. Piccolo S, Dupont S, Cordenonsi M. The biology of YAP/TAZ: hippo signaling and beyond. Physiol Rev. 2014;94(4):1287–312. Epub 2014/10/08. doi: 10.1152/physrev.00005.2014. PubMed PMID: 25287865.

28. Yu FX, Zhao B, Guan KL. Hippo Pathway in Organ Size Control, Tissue Homeostasis, and Cancer. Cell. 2015;163(4):811–28. Epub 2015/11/07. doi: 10.1016/j.cell.2015.10.044. PubMed PMID: 26544935; PubMed Central PMCID: PMCPMC4638384.

29. Yu FX, Guan KL. The Hippo pathway: regulators and regulations. Genes Dev. 2013;27(4):355–71. Epub 2013/02/23. doi: 10.1101/gad.210773.112. PubMed PMID: 23431053; PubMed Central PMCID: PMCPMC3589553.

30. Xin M, Kim Y, Sutherland LB, Qi X, McAnally J, Schwartz RJ, et al. Regulation of insulin-like growth factor signaling by Yap governs cardiomyocyte proliferation and embryonic heart size. Sci Signal. 2011;4(196):ra70. Epub 2011/10/27. doi: 10.1126/scisignal.2002278. PubMed PMID: 22028467; PubMed Central PMCID: PMCPMC3440872.

